# Characterization of Cancer Stem Cells in Laryngeal Squamous Cell Carcinoma by Single-Cell RNA Sequencing

**DOI:** 10.1101/2024.01.21.576534

**Authors:** Yanguo Li, Chen Lin, Yidian Chu, Zhengyu Wei, Qi Ding, Shanshan Gu, Hongxia Deng, Qi Liao, Zhisen Shen

## Abstract

Cancer stem cells (CSCs) constitute a pivotal element within the tumor microenvironment (TME), driving the initiation and progression of cancer. However, the identification of CSCs and their underlying molecular mechanisms in laryngeal squamous cell carcinoma (LSCC) remains a formidable challenge. We employed single-cell RNA sequencing of matched primary tumor tissues, paracancerous tissues, and local lymph nodes from three LSCC patients. Two distinct clusters of stem cells originating from epithelial populations were delineated and verified as CSCs and normal stem cells (NSCs) respectively. CSCs were abundant in the paracancerous tissues compared to the tumor tissues. CSCs showed high expression of stem cell marker genes such as *PROM1*, *ALDH1A1*, and *SOX4*, and increased activity of tumor-related hypoxia, Wnt/β-catenin, and notch signaling pathways. We then explored the intricate crosstalk between CSCs and the TME cells and identified targets within the TME that related with CSCs. We also find eight marker genes of CSCs that correlated significantly with the prognosis of LSCC patients. Furthermore, bioinformatics analyses showed that drugs such as erlotinib, OSI-027, and ibrutinib selectively targeted the CSC-specifically expressed genes. In conclusion, our results represent the first comprehensive characterization of CSCs properties in LSCC at the single-cell level.

## Introduction

Laryngeal carcinoma is the eleventh most common malignancy worldwide and one of the most common type of head and neck cancer. Laryngeal squamous cell carcinoma (LSCC) accounts for 85−95% of laryngeal carcinoma cases [1]. In recent years, improved treatments have increased the survival rates of patients with laryngeal carcinoma, but the recurrence or metastasis rates are still high at 30−40% [2]. Furthermore, aggressive therapies adversely affect phonation, respiration, and deglutition in the LSCC patients and are associated with reduced quality of life and survival [3]. Therefore, there is an urgent need to better understand the mechanisms of tumor invasion, relapse after surgery, and metastasis in LSCC.

According to the cancer stem cell (CSC) theory, a small population of CSCs endowed with properties of self-renewal, tumorigenic, and multi-lineage differentiation are hidden in the cancer tissues and drive tumor progression, therapeutic resistance, and recurrence [4–6]. The development of CSCs was regulated by several transcription factors, intracellular signaling pathways and extracellular niches, such as *SOX2*, notch signaling pathways, and hypoxia, which have been considered as therapy targets to inhibit the biological activities of CSCs [7]. In previous study, scientists analyzed Hep2 and TU-177 LSCC cell lines and identified *PROM1*^+^ *CD44*^+^ cells as the CSCs [8]. However, in tumor tissues, the proportion of CSCs is low and estimated to be about 0.01−2% [7]. Therefore, it is highly challenging to isolate the CSCs from the tumor tissues.

In recent years, single-cell RNA sequencing (scRNA-seq) has been widely used to investigate the regulation mechanisms of tumor heterogeneity, tumor cell subpopulations, and tumor drug resistance [9, 10]. ScRNA-seq has also been shown as a useful technique for studying the characteristics, functions, and molecular mechanisms of the CSCs. For example, scRNA-seq analysis of hepatocellular carcinoma cell lines and tissues showed CSCs characteristics at the single-cell level and identified the marker genes of CSCs with prognostic significance in patients with hepatocellular carcinoma [11]. In addition, scRNA-seq in colorectal cancer showed rare CSCs exist in dormant state and display plasticity toward cancer epithelial cells [12]. Moreover, the marker genes of CSCs were also associated with prognosis of patients with colorectal cancer [12]. These studies have not only underscored the critical role of CSCs in determining cancer prognosis but have also demonstrated the significant advantages of scRNA-seq in identifying key marker genes and unraveling the molecular mechanisms underlying CSCs. However, the distribution and function of CSCs in the pathological tissues of LSCC have not yet been studied at the single-cell level.

In this study, scRNA-seq analysis of matched primary tumor tissues, paracancerous tissues, and local lymph nodes from LSCC patients was performed to comprehensively characterize the CSCs in LSCC. We analyzed the copy number variation (CNV) and marker genes expression in epithelial cell subpopulations, identifying a subpopulation with stem cell characteristics that exist in the early stage of epithelial development. Utilizing immunohistochemistry, function enrichment analysis, transcription factor activity analysis, and validation with public datasets, we were able to classify stem cells into CSCs and normal stem cells (NSCs), respectively. Ligand-receptor analysis revealed extensive communication between CSCs and tumor microenvironment (TME). Furthermore, the marker genes of CSCs were found to correlate with prognosis of LSCC patients, and the potential drugs targeting specifically expressed genes of CSCs were identified. Our results enhance the understanding of how CSCs contribute to tumor progression and offer novel therapeutic strategies for LSCC.

## Results

### Cellular landscape of LSCC

Based on the preceding data from our research group [13], we included nine samples, comprising matched LSCC tissues (LC), paracancerous tissues (PT), and local lymph nodes with tumor metastasis (LM) from three patients diagnosed with LSCC. The behavioral and clinicopathological details of the study subjects are shown in Table S1. Single-cell suspensions were prepared from fresh tissue samples and subjected to scRNA-seq analyses (**Figure 1A**). After applying quality control measures, we obtained 14,748 cells from LC, 17,191 cells from LM, and 8365 cells from PT (Figure S1A–B). These high-quality cells were further assigned into eight cell types according to marker genes expression, including B cells, endothelial cells, epithelial cells, fibroblasts, myeloid cells, T cells, plasmacytoid dendritic cells (pDC), and mast cells (**Figure 1B**, Figure S1C). We then compared the proportions of each cell type in LSCC and observed that T cells were one of the most abundant immune cell type across all specimens (**Figure 1C**), which was consistent with the study conducted by Sun et al [14]. B cells were abundant in LM and accounted for 34.02% of the total cells; however, few infiltrations were observed in LC and PT (Figure S1D). Myeloid cells were enriched in LC (23.83%) but were less abundant in LM and PT (< 10%, Figure S1d), suggesting an inflammatory phenotype in LC.

**Figure 1.**
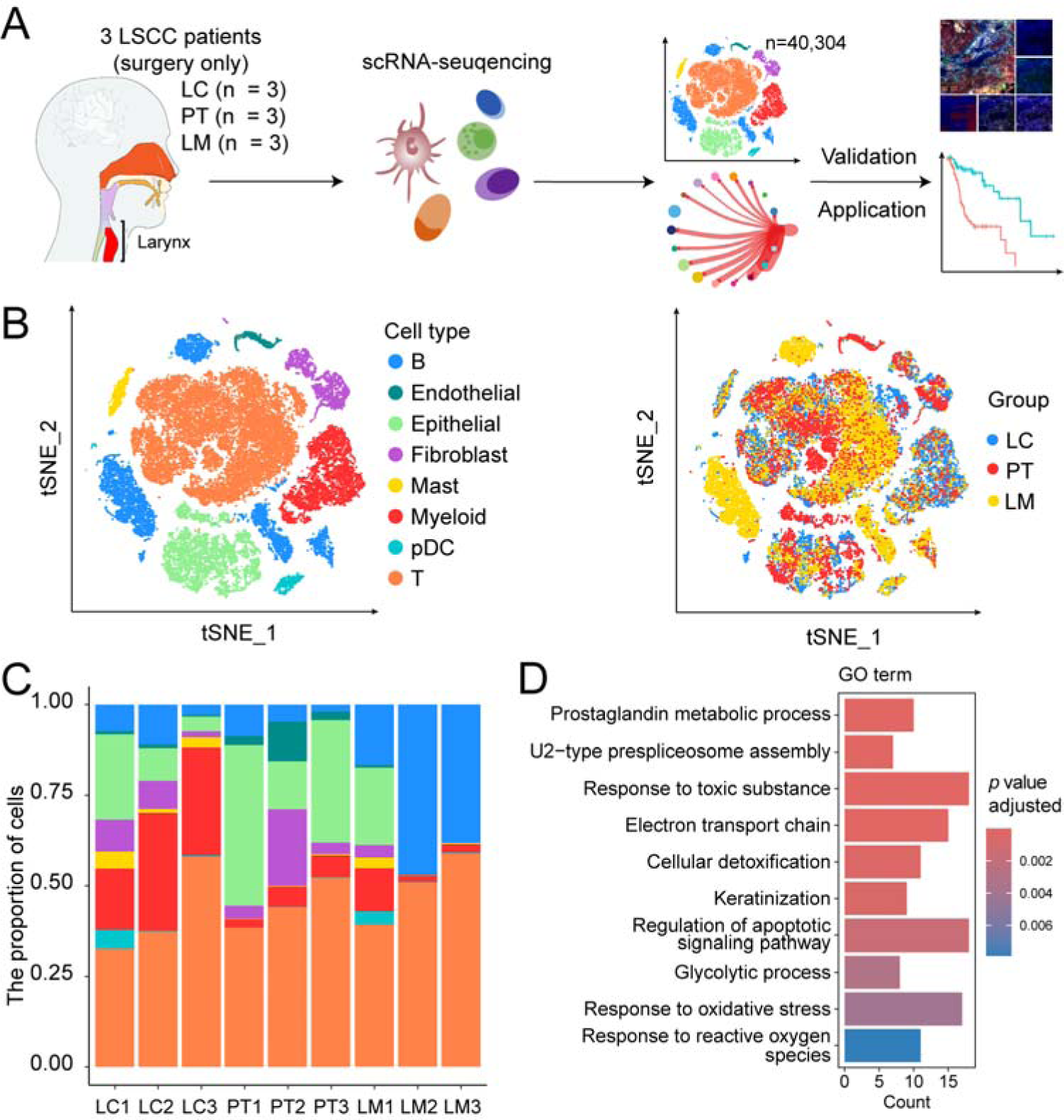
Single-cell transcription profiles of matched tumor tissues, paracancerous tissues, and local lymph nodes from LSCC patients. **A.** Overview of the experimental design. **B.** The t-SNE plot shows categorization of 40,304 single cells into 8 major cell types. The cell types (left) and tissue types (right) are color-coded. Abbreviations: B, B cell; pDC, plasmacytoid dendritic cell; T, T cell; LC, LSCC tissues; PT, paracancerous tissues; LM, local lymph nodes. **C.** Bar plot shows the proportion of different cell types in each sample, colored by the cell types. **D.** GO enrichment analysis based on marker genes of epithelial cells in LC compared to those in PT.

The clinical phenotypes of the three LSCC patients were distinct based on the infiltration patterns of epithelial cells. Patient No. 3 showed the smallest and poorly differentiated tumor lesion compared to other patients, with the highest number of infiltrating T cells and notably fewer tumor epithelial cells in both LC and LM (Figure 1C, Table S1). Next, we conducted a comparative analysis of marker gene expression in epithelial cells between LC and PT. Epithelial cells of LC exhibited high expression of genes associated with metabolism, glycolytic, and response to oxidative stress (**Figure 1D**), including *AKR1C1*, *LDHA*, and *NTRK2* (Table S2). Among them, *LDHA* plays a key role in the Warburg effect or aerobic glycolysis, which is commonly observed in cancer cells under hypoxic conditions [15]. However, the highly expressed genes in epithelial cells from PT tissue such as *KRT4*, *SPINK5*, *CD74*, and *NR4A1* are involved in pathways that regulate antimicrobial humoral response, antigen processing and presentation, and cell-cell adhesion (Figure S1E). Our analysis contributes to a detailed depiction of the TME in LSCC, laying the groundwork for understanding of LSCC.

### Heterogeneity of epithelial lineages and identification of stem cells in LSCC

To classify and delineate the heterogeneity of epithelial lineages in LSCC, we conducted an integrated re-clustering approach to identify subpopulations of epithelial cells across all samples. Eight classes were discerned and annotated, including stem cells (SC), EP-C1 to EP-C6, and Ciliated cells (**Figure 2A**, Figure S2A). CNV analysis indicated that EP-C1, EP-C2, SC, and ciliated cells were in a non-malignant state and enriched in LC (**Figure 2B**). In contrast, EP-C3, EP-C4, EP-C5, and EP-C6, were identified as malignant cells and found to infiltrated both LC and LM (Figure 2B). Notably, gene set variation analysis (GSVA) of hallmark gene sets [16] demonstrated that pathways related with the hypoxia response, tumorigenesis, inflammatory, and interferon responses were enriched in the EP-C4 subset (**Figure 2C**). In contrast, the other malignant epithelial lineages exhibited high activity in *MYC* signaling pathway but a low activity in immune response pathway (Figure 2C). Cell cycle phase scoring demonstrated that EP-C5 and Ep-C6 subsets were in S/G2M phase (Figure S2B). These results were consistent with the high activity of G2M checkpoint and E2F targets signaling pathways (Figure 2C). Since epithelial mesenchymal transition (EMT) promotes metastasis in head and neck cancer, we applied GSVA to analyze the enrichment of EMT gene set across all the epithelial cell subsets [17]. The results showed that the EMT enrichment score was high only for the Ep-C3, EP-C4, and Ep-C5 subsets in the LC tissues of patient No. 3 (Figure S2C). This suggested a patient-specific occurrence of EMT.

**Figure 2.**
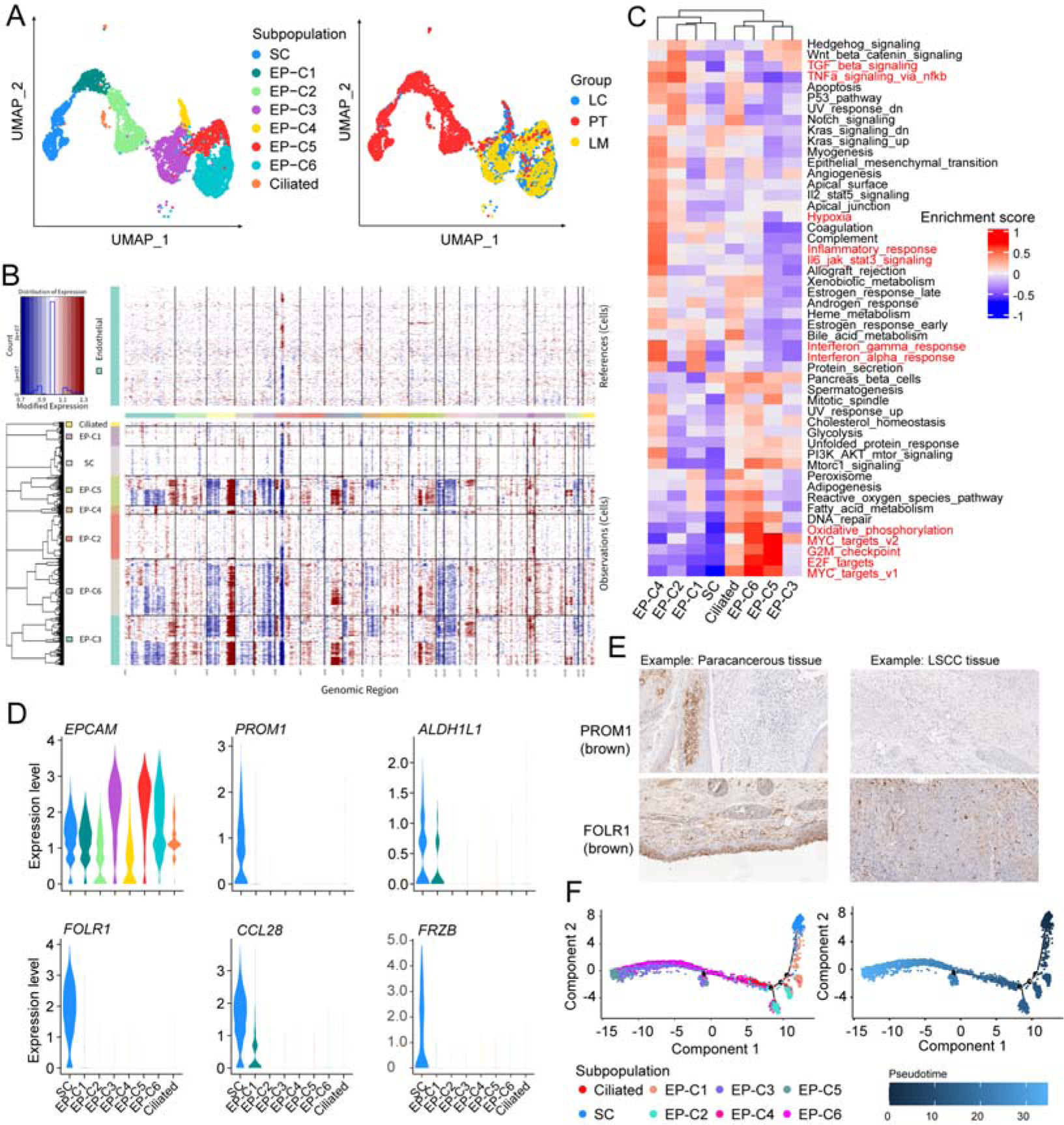
Heterogeneity of epithelial cells and identification of stem cells in LSCC. **A.** UMAP plot shows the heterogeneity of epithelial cells across all the LSCC samples. The epithelial cells were assigned into eight subpopulations, including stem cells (SC), epithelial cells cluster 1 (EP-C1), EP-C2, EP-C3, EP-C4, EP-C5, EP-C6, and ciliated cells (left). The epithelial cell subpopulations (left) and groups (right) are color-coded. **B.** The estimation of the copy number variation (CNV) in the eight epithelial cell subpopulations based on inferCNV. Endothelial cells were used as reference cells. **C.** GSVA of the hallmark gene sets shows activities of different pathways in the eight epithelial cell subpopulations. **D.** Gene expression of the eight epithelial cell subpopulations. **E.** Immunofluorescence staining of *PROM1* and *FOLR1* in the paraffin embedded LSCC and paracancerous tissues. The proteins detected by the corresponding antibodies are indicated on the left. **F.** Pseudotime trajectories show the evolutionary process of epithelial subpopulations in LSCC.

Using the conventional marker genes of *PROM1* and *ALDH1A1* [18], we identified a rare subpopulations of SC (n = 691) (**Figure 2D**). Furthermore, the comparative gene expression analysis between SCs and other epithelial cells showed other novel marker genes for SC, including *FOLR1*, *DMBT1*, *CCL28*, and *FRZB* (Figure 2D, Table S3). Notably, 92.62% and 7.38% of SC were localized in the PT and LC tissues respectively, but none were found in the LM tissues (Figure 2A). Immunofluorescence staining analyses corroborated these findings and demonstrated higher expression levels of *PROM1* and *FOLR1* in paracancerous tissue than in tumor tissues (**Figure 2E**). Moreover, 98.26% of the SCs were in the G1 phase and the gene expression of the cell cycle-related genes in the SCs was significantly low (Figure S2A–B). This further supported the prevailing notion that SCs generally maintain a quiescent or slow-cycling state compared to cancer cells [18, 19]. GSVA results also showed that the expression levels of genes involved in DNA repair, oxidative phosphorylation, and other metabolic pathways were comparatively lower in the SCs compared to the cancer cells (Figure 2C), further corroborating the quiescent state of the SCs. Trajectory analysis demonstrated that the epithelial cells originated from the SCs and branched sequentially towards the non-malignant cells and the malignant cells (**Figure 2F**), demonstrating the high differentiation ability of the SCs.

### Discrimination and genetic properties of cancer stem cell in LSCC

Tumor tissues may concurrently harbor both CSCs and NSCs [20]. Therefore, we performed dimensional reduction and clustering analysis of 691 SCs and categorized them into two distinct clusters, SC-C1 and SC-C2 (**Figure 3A**). This stratification enabled us to ascertain whether the cells exhibited characteristics indicative of CSCs or NSCs. Notably, there were significant differences in the distribution of SC-C1 and SC-C2 between tissues. About 94.12% of SCs in the LC and 32.19% of SCs in the PT were categorized as SC-C2, whereas 5.88% of SCs in the LC and 67.81% of SCs in the PT were categorized as SC-C1 (fisher’s exact test *p* value < 2.2E−16, **Figure 3B**). However, SCs were absent in the LM tissues, which may align with their lower capabilities for EMT feature (Figure S2C). Further analyses of marker gene expression between the two clusters demonstrated upregulation of 541 genes in SC-C2, including *PROM1*, *DMBT1*, *SOX4*, and *FOXQ1* (**Figure 3C**, Table S3). Gene Ontology (GO) enrichment analysis showed that these 541 genes were significantly associated with tumor-related pathways, including cell adhesion, response to tumor necrosis factor, response to oxidative stress, and apoptotic signaling (Figure S2D). In contrast, 85 genes were upregulated in SC-C1, and these genes were not associated with any tumorigenesis-related pathways (Figure S2E).

**Figure 3.**
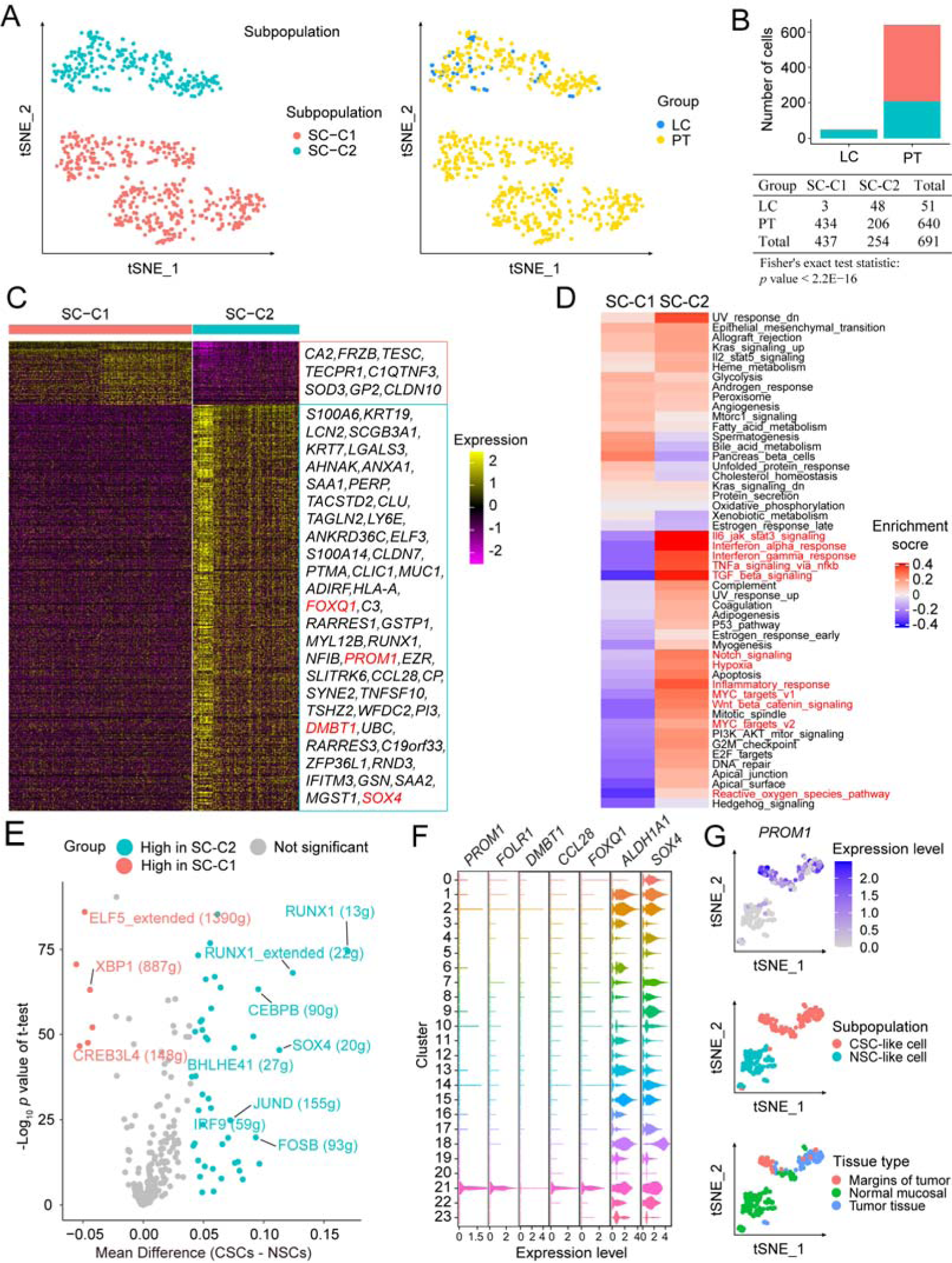
Identification and characterization of cancer stem cells in LSCC. **A.** The t-SNE plot shows the distribution of two epithelial stem cell cluster in LSCC. **B.** Bar plot shows absolute number of two stem cell subpopulations. Fisher’s exact test was used to assess the statistical significance between groups. **C.** Heatmap shows expression levels of the marker genes in the two stem cell subpopulations (Table S3). **D.** GSVA analysis of the hallmarker pathway activities between SC-C1 and SC-C2. **E.** The volcano plot shows mean differences in the AUC scores for each regulon between the cancer stem cells (CSCs) and normal stem cells (NSCs). *P* values were calculated using the student’s t-test. **F**. The expression of CSC marker genes in an independent validation scRNA-seq dataset (GES206332). **G**. Re-clustering of cluster 21 from Figure 4F revealed that CSC-like cells highly expressed *PROM1* and are distributed across various tissue types.

Subsequently, enrichment analysis of hallmark gene sets demonstrated that SC-C2 displayed heightened activity of Wnt/β-catenin signaling, notch signaling, hypoxia, MYC targets, inflammatory, and interferon response pathway (**Figure 3D**). Conversely, most of these pathways were inactive in SC-C1 (Figure 3D). A previous study by Sato et al demonstrated that the expression of Wnt-3 and Notch-ligand on the *CD24*^+^ Paneth cells was necessary for the stemness maintenance of the *LGR5*^+^ stem cells in colon crypts [21]. Therefore, we hypothesized that activation of Wnt/β-catenin and notch signaling pathways contributed to the maintenance of stemness in the SC-C2 population. Furthermore, we analyzed the transcription factor activity using the single-cell regulatory network inference and clustering (SCENIC) method [22] and estimated the mean differences in the area under the curve (AUC) scores of all the regulons (transcription factor and its target genes) between the SC-C1 and the SC-C2. SCENIC analysis results showed higher AUC scores for the *RUNX1* (13g) and *SOX4* (20g) regulons in the SC-C2 compared to SC-C1 (**Figure 3E**). In previous study, *RUNX1* was found to collaborate with the androgen receptor in triple-negative breast cancer CSCs to promote disease recurrence after chemotherapy, whereas inhibition of *RUNX1* transcriptional activity reduced expression of CSC marker genes [23]. Aberrant overexpression of *SOX4* is also associated with the development of multiple human cancers and the maintenance of cancer cell stemness [24].

In summary, given the unique characteristic of SC-C2 cells, they may function as CSCs, while SC-C1 cells appear to serve as NSCs. We then focused on validating CSC using public dataset. We involved a reanalysis of epithelial cells defined by Sun et al [14] through scRNA-seq data collected from three LSCC patients (GES206332). We found that cluster 21 (n = 220) strongly and specifically expressed the SC marker genes identified in our analysis (**Figure 3F**). Further re-clustering of these 220 cells revealed one cluster expressing the marker genes of CSCs, which were named as CSC-like cells, while another cluster were named as NSC-like cells (**Figure 3G**). Additionally, we observed the distribution of CSC-like cells in tumor tissue, margins of tumor, and normal mucosal adjacent to the tumor of LSCC (**Figure 3F**, Figure S2F). Our results thus provide robust evidence and a comprehensive characterization of identified CSCs in LSCC.

### Clinical evaluation of cancer stem cells signatures in prognosis and potential drug responses

Given the crucial role of CSCs (Cancer Stem Cells) in cancer development, recurrence, and metastasis, the marker genes of CSCs have significant potential for clinical applications. Therefore, we screened the marker genes by comparing the gene expression patterns between the CSCs and all the other cell types using the fraction of gene detection > 0.3, adjusted *p* value < 0.01, and average log_2_(fold change (FC)) > 0.5 as threshold parameters. As a result, 172 CSC marker genes, such as *FOLR1*, *PROM1*, *SAA2*, *DMBT1*, and *CCL28*, were identified (**Figure 4A**, Table S4). GO enrichment analysis demonstrated that the CSC marker genes were associated with epidermis development, negative regulation of hydrolase activity and proteolysis, and endopeptidase inhibitor activity (Figure S3A). Multiplex immunofluorescence confirmed the co-localization of *PROM1*, *FOLR1*, and *DMBT1* in epithelial cells, which were positively identified by cytokeratin (CK) (**Figure 4B**). Results indicating these CSC marker genes can be considered as biomarkers to discriminate the CSCs in LSCC.

**Figure 4.**
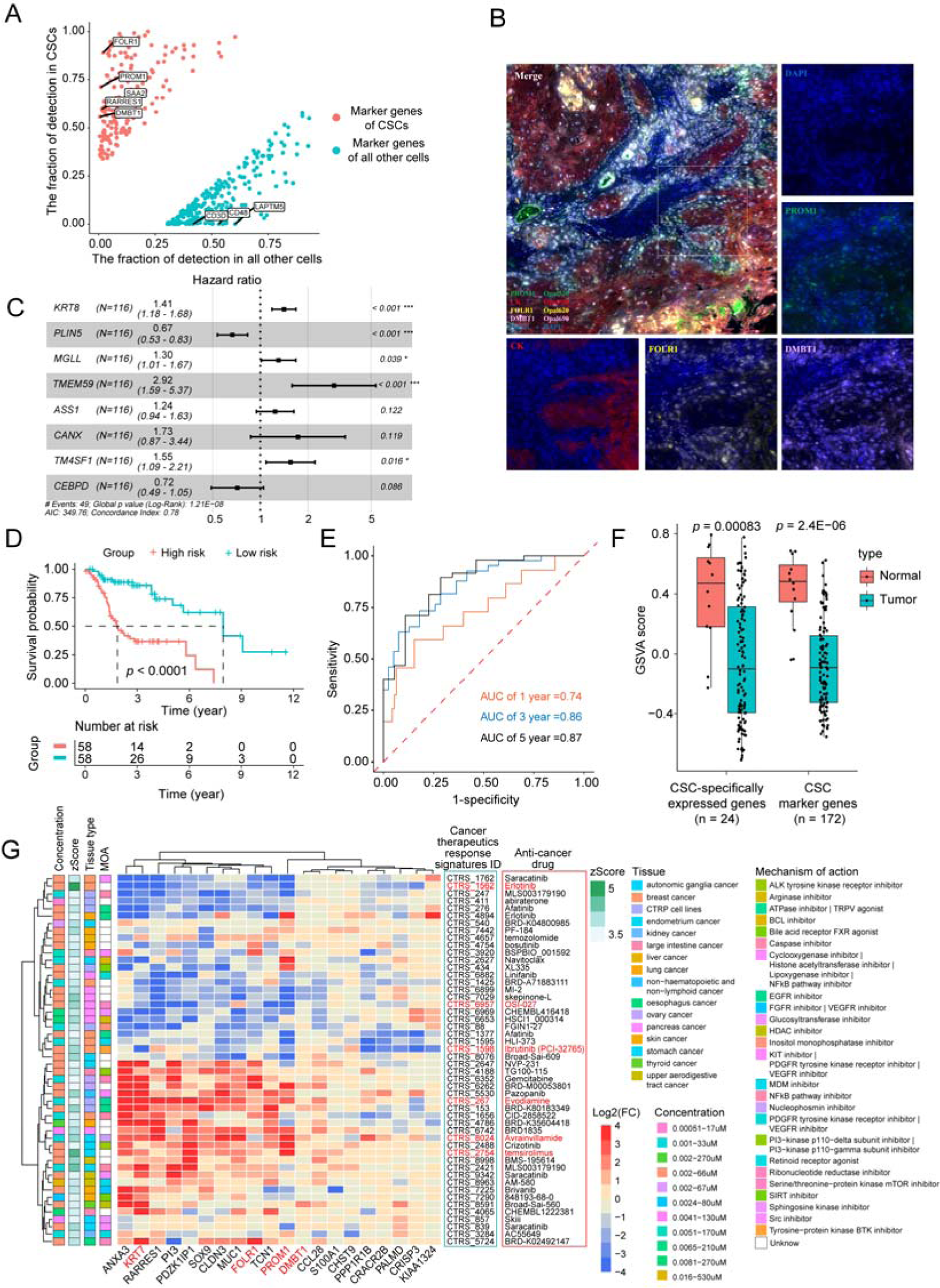
Clinical application of CSC signatures. **A.** Point plot shows fraction of marker genes detected in CSCs compared with all the other cell types (Table S4)**. B.** Confocal multiple immunofluorescence images shows the expression levels of cytokeratin (CK), *PROM1*, *DMBT1*, and *FORL1* proteins in the LSCC tissues. **C.** Forest plot shows the hazard ratio of prognosis signatures in the model based on eight CSC marker genes**. D.** Kaplan-Meier survival curves show the survival rates of the high-risk and low-risk group of LSCC patients based on the eight-gene prognosis signature. **E.** ROC curves show the 1-, 3- and 5-year overall survival of the LSCC patients as predicted by the risk score calculated using the eight-gene prognosis signature. **F.** Bar plot shows the GSVA scores for the CSCs signatures in 116 LSCC samples from TCGA. **G.** Heatmap shows the expression pattern of query genes in Cancer Therapeutics Response Signatures. Genes with negative values were associated with drug sensitivity, whereas genes with positive values were associated with drug resistance. The term “mechanism of action” refers to specific biochemical interaction by which a drug produces pharmacological effects.

Subsequently, we performed LASSO and Cox regression analyses to establish a prognostic prediction model based on the CSC marker genes using 116 LSCC samples from The Cancer Genome Atlas database (TCGA, Figure S3B). The Cox proportional hazard regression model with eight CSC marker genes (*KRT8*, *PLIN5*, *MGLL*, *TMEM59*, *ASS1*, *CANX*, *TM4SF1*, and *CEBPD*) was constructed to calculate the risk scores for the LSCC patients and divided them into low-risk and high-risk groups using the median risk score (**Figure 4C**). Kaplan-Meier survival curves demonstrated that the survival rates were higher for the low-risk patients compared with the high-risk group (**Figure 4D**). Time-dependent ROC analysis demonstrated satisfactory prognostic prediction accuracy, with AUC values of 0.74, 0.86, and 0.87 for the 1-, 3-, and 5-year survival rates, respectively (**Figure 4E**). Furthermore, we validated the model accuracy using the TCGA dataset by dividing the patients into training (70%) and testing (30%) datasets. Time-dependent ROC analysis showed that the AUC values were higher than 0.8 for both 3- and 5-year survival in both the training and the testing datasets (Figure S3C). Therefore, our prognostic model proves to be a valuable tool in predicting the outcomes for patients with LSCC.

We also identified genes specifically expressed in the CSCs from marker genes by using the rigorous threshold criteria with the fraction of gene detection in CSCs > 0.5 and < 0.05 in all other cells. Consequently, 24 genes, including *FOLR1*, *PROM1*, *DMBT1*, and *SOX9*, were identified as CSC-specifically expressed genes (Table S4). GSVA results showed that these CSC-specifically expressed genes were significantly enriched in the paracancerous samples compared with the tumor samples from TCGA LSCC dataset (**Figure 4F**) and were consistent with our findings above. Then, the relationships of these CSC-specifically expressed genes with the pre-computed Cancer Therapeutics Response Signatures were assessed using iLINCS tool [25, 26]. The results showed that several potential anti-cancer drugs, including erlotinib, OSI-027, and ibrutinib (PCI-32765) repressed CSCs by inhibiting the expression of CSC-specifically expressed genes (**Figure 4G**, Table S5). Our data also showed that the drugs such as temsirolimus, avrainvillamide, and evodiamine, induced resistance by upregulating these CSC-specifically expressed genes (Figure 4G, Table S5). Our findings identified potential drugs that can be used to selectively suppress the CSCs without affecting NSCs and other cell types.

### Crosstalk between tumor immune microenvironment and cancer stem cells in LSCC

Elucidating the interplay between CSCs and their ecological niches is critical for understanding the regulation mechanisms of cancer development and metastasis, as well as the identification of novel therapeutic targets in the TME. We subsequently performed unsupervised clustering of the major types of immune cell, including T cells, myeloid cells, and B cells. Following this, we investigated the intricate cell-cell communication network between the CSCs and distinct subpopulations of immune cell.

T cells were the predominant immune cell type with seven distinct subpopulations based on the expression levels of the conventional marker genes (**Figure 5A**, Figure S4A). The proportion of regulatory T cells (Treg-FOXP3) was higher in the LC and LM tissues than in the PT (Figure 5A), suggesting the presence of an immunosuppressive microenvironment in the LC and LM tissues. Furthermore, two clusters of CD8^+^ T cells demonstrated exhaustion because of higher expression levels of exhausted markers (*PDCD1*, *LAG3*, and *TIGIT*) and were categorized as exhausted cells (Figure S4A). Myeloid cells were categorized into four clusters (**Figure 5B**, Figure S4B). Among them, TAM-APOE, which represented the tumor-associated macrophages with high expression levels of *APOE*, *CD163*, and *VSIG4* (Figure S4B). Besides, four discrete subpopulations of B cells were identified, including follicular B cells, memory B cells, plasma cells, and marginal zone B cells (**Figure 5C**, Figure S4C).

**Figure 5.**
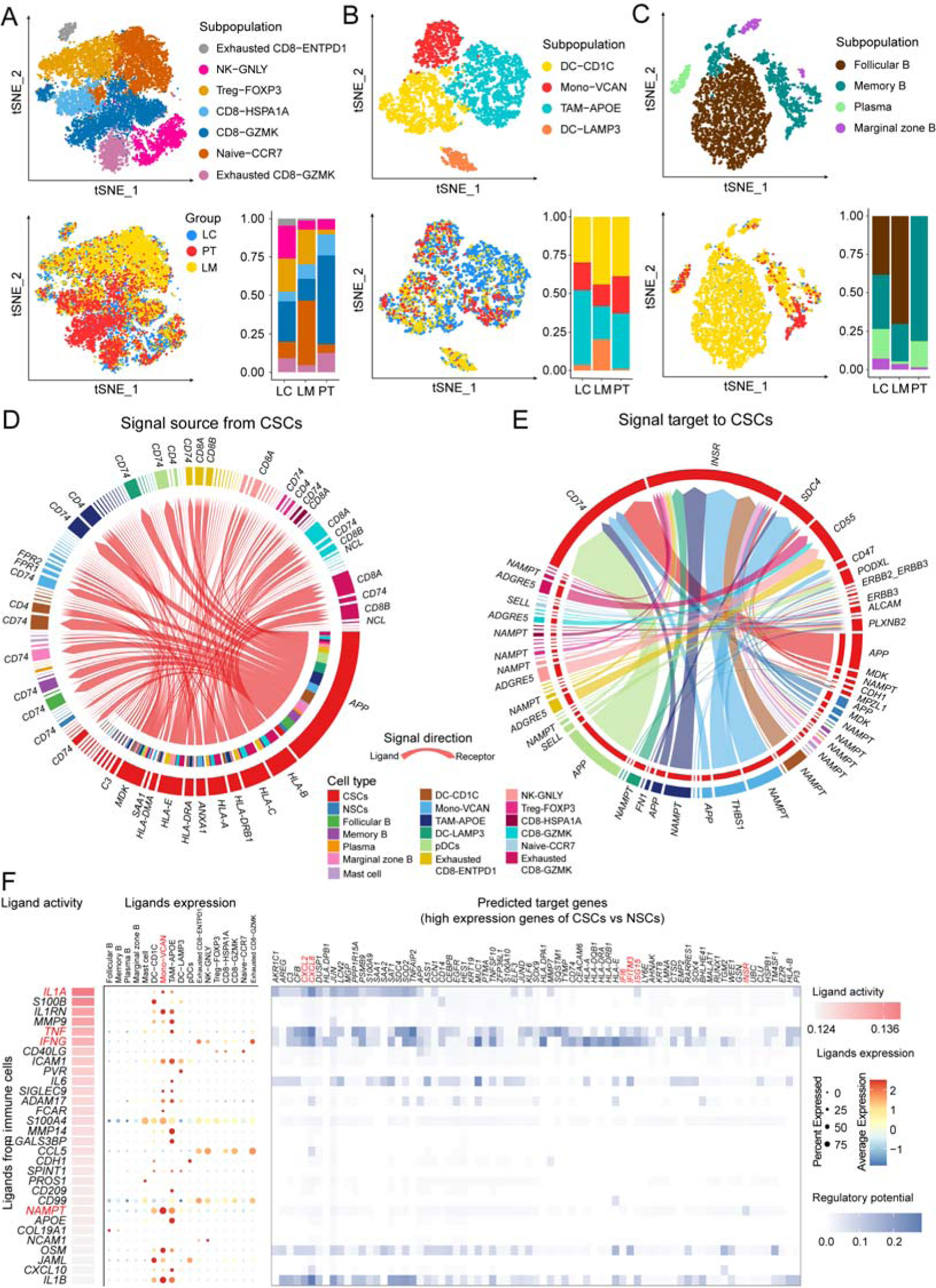
Cell-cell communication between immune cells and cancer stem cells. **A**–**C.** The t-SNE plots of seven T cells subpopulations, four myeloid cell subpopulations, and four B cells subpopulations in all samples. The bar plot shows proportions of various immune cell subpopulations in LC, LM, and PT. The cells are color-coded according to subpopulations (top) and tissues (bottom). **D.** Circle plot shows signaling molecules from CSCs interacting with the immune cells. The edge width denotes the strength of communication. The circles of distinct colors represent distinct cell types. **E.** Circle plot shows signaling molecules from immune cells interacting with the CSCs. The edge width denotes strength of the communication. The circles of different colors represent distinct cell types. **F.** NicheNet analysis shows potential ligands expressed by immune cells that presumably affected the differentially expressed genes between CSCs and NSCs. Ligand activity indicates the ability of each ligand to predict the target genes. Ligands with better prediction are ranked higher. The regulatory potential score indicates the confidence that a particular ligand can regulate the expression of a particular target gene.

Subsequently, we used the CellChat [27] and NicheNet [28] tools to predict the key signaling events between the immune system and the CSCs. Our findings revealed that the CSCs engaged in reciprocal signaling through ligand-receptor interactions with the immune system via 42 outgoing signals and 19 incoming signals (**Figure 5D–E**, Figure S4D–E). The interaction between *APP* and *HLA* was the strongest outgoing signaling interaction between the CSCs and the immune cells (Figure 5D). While the interaction between *INSR* (specifically expressed on the CSCs) and *NAMPT* (expressed on various myeloid lineages) was the strongest incoming signal (Figure 5E). NicheNet analysis also revealed that *NAMPT* induced higher expression of genes in the CSCs compared to NSCs (**Figure 5F**). According to the previous reports, *NAMPT* is a key enzyme in NAD^+^ biogenesis and has been linked to induction of cancer stem cell-like properties in colon cancer [29]. Additionally, *NAMPT* inhibitors suppress senescence-associated CSCs induced by platinum-based chemotherapy in ovarian cancer [30]. Therefore, we postulated that inhibition of *NAMPT* may abrogate the development of CSCs in LSCC. Additionally, NicheNet also highlighted high activities of the ligands such as *IL1A*, *TNF*, and *IFNG* (Figure 5F). This further demonstrated elevated expression of genes related with the inflammatory and interferon responses in the CSCs that described above. These results provided a comprehensive insight regarding the interactions between the CSCs and the immune microenvironment in LSCC.

### Crosstalk between stromal cells and cancer stem cells in LSCC

Endothelial cells and fibroblasts play crucial roles in the development of both cancer cells and the CSCs. In this study, four subpopulations of endothelial cells in LSCC were identified (**Figure 6A**, Figure S4F). The vECs-RGS5 subpopulation exhibited high expression levels of *RGS5* and *EPAS1*, thereby suggesting their role in response to the changes in oxygen levels and angiogenesis [31] (Figure S4F). The vEC-FLT1 subpopulation represented tumor-associated endothelial cells, which were characterized by high expression level of VEGFR1 (*FLT1*); moreover, significant number of vEC-FLT1 endothelial cells were found in the LC tissues (Figure 6A, Figure S4F). We also identified five subpopulations of fibroblasts, Fib-C1 to Fib-C5 (**Figure 6B**). The Fib-C2 cells were predominantly found in the LC and LM tissues and showed high expression levels of mesenchymal-related genes, including *FAP*, *WNT5A*, *POSTN*, and *MME* (Figure 6B, Figure S4G). Therefore, Fib-C2 cells were identified as the cancer-associated fibroblasts [32]. Fib-C3 cells were similar to the vascular cancer-associated fibroblasts (vCAFs) in the intrahepatic cholangiocarcinoma [33] and exhibited high expression levels of [-SMA, *RGS5*, *MYH11* and *EPAS1* (Figure S4G). These results highlighted the characteristics of stromal cells in the proximity of the CSCs.

**Figure 6.**
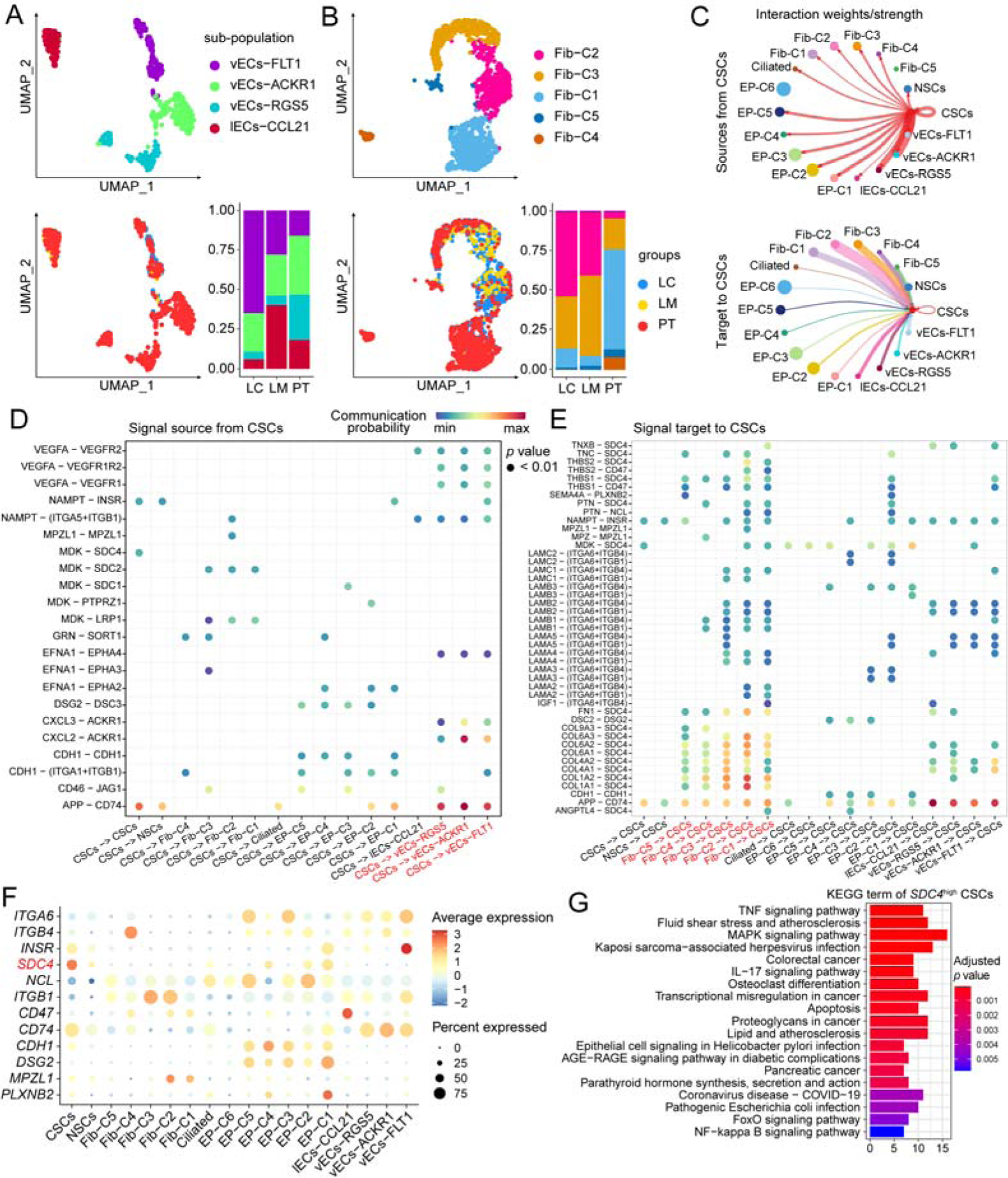
Intercellular communication between stromal cells and cancer stem cells in LSCC. **A**–**B.** UMAP plots show the four endothelial cell subpopulations and five fibroblast subpopulations in all the samples. The bar plot shows proportion of each subpopulation in different tissues. The various subpopulations (top) and tissues (bottom) are depicted by distinct colors. **C.** Circle plot shows signaling molecules from the CSCs that interact with stromal cells and the signaling molecules from stromal cells that interact with the CSCs. The edge width indicates strength of the communication. **D**–**E.** Dot plot shows the ligand-receptor pairs between CSCs and stromal cells. Dot color represents the calculated communication probability; dot size represents the *p* values. **F.** Dot plot shows expression levels of key ligands or receptors in LSCC. **G.** KEGG pathway enrichment analysis results for the marker genes in the *SDC4*^high^ CSCs.

We then performed analysis of the cell-cell communications between the CSCs and stromal cells. The results showed that CSC-related signals primarily targeted the vECs (**Figure 6C**) and involved ligand-receptor pairs such as *VEGFA*-*VEGFR*, *CXCL2*-*ACKR1*, and *APP*-*CD74* (**Figure 6D**). Interestingly, signals targeting the CSCs predominantly originated from fibroblasts, especially Fib-C2 cells (Figure 6C). Based on systematic investigation of the predicted signals from the fibroblasts to the CSCs, we identified 37 ligand-receptor pairs that contributed to ECM-receptor interactions (**Figure 6E**). Among these interactions, the highest probability of communication was observed between collagen from Fib-C2 cells (*COL1A1* and *COL1A2*) and *SDC4* from CSCs (Figure 6E). Moreover, *SDC4*, one of the CSC marker gene, was highly and specifically expressed in CSCs (**Figure 6F**). In previous study, *SDC4* has shown to plays a pivotal role in cell adhesion and migration and is a promising therapeutic target in hepatocellular carcinoma [34]. Besides, Chen et al demonstrated that *SDC4* overexpression promoted EMT and inhibited cancer cell apoptosis by activating the Wnt/β-catenin signaling pathway in human papillary thyroid cancer [35]. This suggests that *SDC4* could serve as potential therapeutic target for addressing CSCs in LSCC. We then further classified CSCs into two groups based on the median expression levels of *SDC4* and performed functional enrichment analyses on differentially expressed genes between the two groups*. SDC4*^high^ CSCs showed high expression of genes such as *KLF6*, *ATF3*, and *CXCL2* (Figure S4H). Kyoto Encyclopedia of Genes and Genomes (KEGG) enrichment analysis demonstrated that TNF signaling, MAPK signaling, apoptosis, and cancer development were involved (**Figure 6G**). Above all, our results elucidated key patterns in the intercellular communication between stromal cells and CSCs and identified potential therapeutic targets for LSCC.

## Discussion

A comprehensive understanding of the identity and characteristics of CSCs in LSCC is critical for enhancing the efficacy of cancer therapy to mitigate cancer recurrence. In this study, we delineated the cellular heterogeneity in LSCC at a single-cell level in matched tumor tissues, paracancerous tissues, and local lymph nodes. Then, both bioinformatic and experimental approaches provided reliable identification and characterization of the CSCs, followed by a comprehensive analyses of marker gene expression, signaling pathway activities, activation of regulons, and molecular interactions with the ecological niche. Furthermore, we screened for the potential prognostic markers and therapeutic targets among the CSC marker genes and CSC-specifically expressed genes in LSCC. In summary, our study sheds new light on the biological function and molecular mechanisms of CSCs in LSCC and offers a valuable resource for the development of CSC-targeted therapeutic strategies for clinical applications.

Several studies have shown that a small population of CSCs can fueled the development and progression of various tumors [4–6]. However, the identification and characterization of these cells remain elusive. We initially analyzed the epithelial stem cells in LSCC by estimating the expression of canonical marker genes, *PROM1* and *ALDH1A1*. Immunofluorescence staining confirmed *PROM1* expression in LSCC, revealing higher levels in the paracancerous tissues compared to the tumor tissues. This suggested the presence of CSCs in both the paracancerous and tumor tissues. We then identified a distinct SC-C2 population with CSC characteristics, especially in LC. SC-C2 cells showed higher *PROM1* and *SOX4* expression, enrichment of tumorigenesis-related and stemness maintenance pathways, and activated regulons associated with the cancer stem cell phenotype. Furthermore, essential signals for stem-cell maintenance, including hypoxia, Wnt/β-catenin, notch, and NF-κB signaling pathways [21] were activated in these CSCs, thereby providing evidence for discriminating the CSCs from other cell types in LSCC. Moreover, several new CSC marker genes we identified in LSCC, including *DMBT1* and *SOX4*. These new CSC marker genes have been previously reported to play significant roles in SCs development [36, 37]. Utilizing public datasets and established CSC marker genes, we confirmed the presence of CSCs in LSCC. Our results showed reliable evidence to distinguish CSCs from other cell types in LSCC.

Previous study has reported CSC gene signature showed good prognostic prediction in the patients with hepatocellular carcinoma [11]. In our study, we constructed a prognosis model from 8 CSCs marker genes, including *KRT8*, *PLIN5*, *MGLL*, *TMEM59*, *ASS1*, *CANX*, *TM4SF1*, and *CEBPD*. The expression levels of these genes correlated with the patient outcomes in LSCC from TCGA. Furthermore, using specific filtering criteria, 24 CSC-specifically expressed genes were identified, including *FOLR1*, *PROM1*, *DMBT1*, and *SOX9*, which showed minimal fraction of gene detection in the other cell types. We also showed that the development of these CSCs in LSCC can be inhibited potentially by targeting the expression of these CSC-specifically expressed genes using small molecule drugs, including erlotinib, OSI-027, and ibrutinib. For example, erlotinib was effective in reducing CSC proliferation, promoting CSC differentiation, and enhancing chemo/radiation treatment sensitivity in head and neck squamous cell carcinoma [38]. Our findings provide valuable insights for designing novel therapeutic strategies to specifically target CSCs in LSCC. Understanding the ecological niche of the CSCs is critical in improving the cancer therapeutic efficacy and identifying novel TME targets for treatment. In this study, we investigated the crosstalk of the CSCs with various immune cell lineages and the stromal cells. The results showed that the interaction between myeloid cells and CSCs through the *NAMPT*-*INSR* signaling pathway significantly affected CSCs. *NAMPT* and *INSR* are potential therapeutic targets in multiple cancer types[29, 30, 39] and may be crucial in suppressing CSCs. Moreover, ligand-receptor interactions such as *VEGFA*-*VEGFR*, *CXCL2*-*ACKR1*, and *APP*-*CD74* promoted angiogenesis via interacting with the endothelial cell receptors, which highlighted the potential role of CSCs in promoting angiogenesis. Additionally, fibroblasts expressed numerous ECM-related molecules, which interacted with the CSCs via *SDC4* and integrins. ECM remodeling is linked with tumor malignancy, survival, and migration [40]. Our results suggested that ECM modeling played a critical role in establishing the niche for the CSCs.

Although many highlights in our study, there are several limitations too. For instance, we successfully identified the characteristics of CSCs and their distinct features compared with NSCs, but we did not investigate the source and evolution of CSCs. Therefore, further investigations are necessary to determine the origin and development of CSCs through *in vitro* isolation, lineage tracing, and genome sequencing in future. In addition, we identified potential therapeutic targets, but further validation and characterization are also required through *in vitro* and *in vivo* experiments.

## Materials and methods

### Patient sample collection and preparation of single cells

This study recruited three LSCC patients that had not received radiotherapy, chemotherapy, or other treatments prior surgery at the Lihuili Hospital of Ningbo University (Ningbo, China). Written informed consent was obtained from all the participants. The ethical approval for this study was obtained from the Institutional Review Board of Lihuili Hospital, Ningbo University (Approval No. KY2020PJ191). After surgery, fresh tissue specimens were immediately cut into small slices, washed two- or three-times using Dulbecco’s phosphate-buffered saline and incubated in a preservation solution at 4 °C. Each tissue sample was cut into slices with approximately 2−4 mm. A Human Tumor Dissociation Kit (Catalog No. 130-095-929, Miltenyi Biotec, Auburn, CA) was then used according to the manufacturer’s manual, and the samples were cultured at 37 °C for 30 min in a digestion solution containing RPMI 1640, Enzyme H, Enzyme R and Enzyme A (Catalog No. 11875093, Thermo Fisher Scientific, Waltham, MA). The cell suspension was strained using a 30-μm cell strainer to remove cell aggregates. The specimens were resuspended in RPMI 1640 and centrifuged at 300 g for 7 min to collect the cell supernatant. DNase treatment was performed on samples that exhibited viscosity after dissociation. Cells were resuspended in an erythrocyte lysis buffer (Catalog No. 130-094-183, Miltenyi Biotec) and processed using a dead cell removal kit (Catalog No. 130-090-101, Miltenyi Biotec) to remove erythrocytes and dead cells, respectively. Cells were stained using 0.4% trypan blue (Catalog No. T10282, Thermo Fisher Scientific) to test viability.

### scRNA-seq and data processing

The sequencing libraries were prepared using the 10x Genomics protocol of cell capture and library construction. The high-quality sequencing libraries were sequenced using the Illumina platform. The sequencing data in the fastq format was aligned to the human reference genome (GRCh38). The processed data was further analyzed using Seurat 4.0 package [41]. The doublets in the scRNA-seq data were detected and handled using the scDblFinder [42]. Subsequently, genes detected in less than one cell were removed, and the cells satisfying the following requirements were retained: > 1000 unique molecular identifier (UMI) counts; > 700 genes; < 20% of mitochondrial gene expression in the UMI counts; < 0.1% hemoglobin gene expression in the UMI counts; and the value for log_10_(genes) divided by log_10_(UMI) was > 0.75.

The gene expression matrices were then normalized using “SCTransform” in Seurat, and the variable genes were selected using a residual variance cutoff value of 2.3. Moreover, mitochondrial genes, ribosomal genes, and cell-cycle genes (ccgenes in Seurat) were removed from variable genes. PCA and harmony algorithm [43] with default parameters were used for batch correction. The cell clusters were identified using the shared nearest neighbor modularity optimization-based clustering algorithm and resolutions between 0.1 to 1.5 were applied to identify better subcluster representation and robustness. The clustering results were visualized with the UMAP and t-SNE projections.

### Cell type assignment and marker gene identification

The major cell types were identified based on the expression and distribution of canonical marker genes and were further verified with SingleR [44]. The marker genes of each cluster were determined using the MAST test [45] using the “FindAllmarker” function with positive average log_2_(FC) higher than 0.5 and adjusted *p* value lower than 0.01 unless mentioned otherwise. The subpopulation of major cell types were also identified by the similar pipeline as described above.

### Trajectory, copy number variations analysis, SCENIC analysis, and cell–cell communication analysis

We applied Monocle2 [46] with default parameters to determine the potential lineage differentiation of epithelial cells. The variable genes were selected by Seurat as mentioned above. We used the inferCNV (v1.2.1) to estimate the CNVs of all the epithelial subpopulations and the endothelial cells were used as reference. To infer gene regulatory networks based on co-expression and DNA motifs, we performed SCENIC analysis of the scRNA-seq data with default parameters. The cell-cell communication analysis was analyzed using CellChat with default parameters. The NicheNet “nichenet_seuratobj_cluster_de” function was used to explain differential expression between two ‘receiver’ cell clusters by ligands expressed by niche cells.

### Function enrichment and gene set activity analysis

GO and KEGG pathway enrichment analyses were performed using the clusterProfiler [47]. GSVA was performed to estimate the activity of the gene sets using default parameters based on the hallmarker gene sets (MsigDB, v7.4). The average expression levels of genes in each subpopulation were used as input for GSVA.

### Construction of CSC marker gene signatures and abundance estimation

The CSC marker genes were defined as those with a positive average log_2_(FC) > 0.5, adjusted *p* value < 0.01, and fraction of detection > 0.3 when compared with all the other cells. The CSC-specifically expressed genes were identified from CSC marker genes based on a fraction of genes detected > 0.5 in the CSCs and < 0.05 in all the other cells. Then, GSVA was used to evaluate the abundance of CSCs in all the LSCC samples from the TCGA database based on the above two gene sets (CSC marker genes and CSC-specifically expressed genes) and higher enrichment scores indicated higher number of CSCs.

### Drug prediction

To identify drugs that can regulate expression levels of the CSC-specifically expressed genes, we first applied iLINCS web server to calculate the connectivity between these genes and the Cancer Therapeutics Response Signatures. Next, the top 50 drugs with highest connectivity were selected for further analysis. Next, the top 50 drugs with highest connectivity were selected for further analysis. To further determine the drug effects, we used morpheus analysis, a web tool of iLINCS, to construct a perturbation matrix for these drugs to the CSC-specifically expressed genes. The perturbation matrix was visualized and clustered in R. Genes with negative values were associated with drug sensitivity, whereas genes with positive values were associated with drug resistance. The mechanism of action was defined as the specific biochemical interaction through which a drug produced its pharmacological effects.

### Survival analysis of the LSCC-TCGA dataset

The expression profiles and clinical characteristics of TCGA-HNSC patients were downloaded from UCSC Xena database. We only included patients with LSCC. LASSO regression and Cox regression analyses were performed in a stepwise manner to establish a prognostic prediction model with the CSC marker genes using the survival and glmnet packages [48]. We first established the prognosis model based on the entire cohort and calculated a risk score for each patient. Then, we divided the entire cohort based on a 7:3 ratio to validate the predictive accuracy of the prognosis model.

### Immunohistochemical and multiple immunofluorescence staining

Immunohistochemical staining for *FOLR1* (Catalog No. PA5-42004, Thermo Fisher Scientific) and *PROM1* (Catalog No. MA1-219, Thermo Fisher Scientific) was performed using the EnVision two-step method [49]. Briefly, the tissue slices were incubated at 100°C with the EDTA antigen repair solution for 20 min. The samples were then cooled to room temperature. The results of immunohistochemical staining were evaluated by two pathologists using a double-blind method.

We also performed multiple immunofluorescences staining with the *DMBT1* (Catalog No. PA5-83517, Thermo Fisher Scientific), *FOLR1* (Catalog No. PA5-42004, Thermo Fisher Scientific), *PROM1* (Catalog No. MA1-219, Thermo Fisher Scientific), CK antibodies, and DAPI. CK-positive cells represented epithelial cells and indicated greater likelihood of epithelial-derived cancer. The opal 7-color Manual IHC Kit (Catalog No. NEL801001KT, PerkinElmer, Shelton, CT) and VECTASHIELD® HardSet Antifade Mounting Medium (Catalog No. H-1400, Vector Laboratories, Newark, CA) were used for immunofluorescences analysis. In contrast to the conventional staining process, the primary and secondary antibodies were cleaned by microwaving after the opal dyes were colored. The samples were then stained with the other antibodies until all labeling was complete.

## Ethical statement

The biospecimens used in this study were provided by the Lihuili Hospital of Ningbo University. Written informed consent was obtained from all participants, and ethical approval was obtained from the Institutional Review Board of Lihuili Hospital of Ningbo University (KY2020PJ191).

## CRediT author statement

**Zhisen Shen and Qi Liao:** Conceptualization, Writing - Review & Editing, Supervision, and Funding acquisition. **Yanguo Li:** Methodology, Software, Data Curation, Formal analysis, Visualization, Writing - Original Draft. **Chen Lin and Yidian Chu:** Resources, Investigation, Data Curation, Validation. **Shanshan Gu, Hongxia Deng, and Zhengyu Wei:** Resources, Investigation. **Qi Ding:** Validation, Investigation.

## Acknowledgements

This research project is supported by Ningbo Top Medical and Health Research Program (Grant No. 2023030514); Ningbo Clinical Research Center for Otolaryngology Head and Neck Disease (Grant No. 2022L005); Ningbo medical and health brand discipline (Grant No. PPXK2018-02); Zhejiang Provincial Natural Science Foundation of China (Grant No. LY19H160014, LQ21H130001); Ningbo “Technology Innovation 2025” Major Special Project (Grant No. 2020Z097); Ningbo Natural Science Foundation (Grant No. 2023J030, 2021J290), all of which are in China.

## Supplementary material

### Supplementary Tables

Table S1 The clinical information and pathologic features of patients

Table S2 The differentially expressed genes of epithelial cells between LC and PT

Table S3 The differentially expressed genes of SC-C1 and SC-C2

Table S4 Cancer stem cell gene signatures

Table S5 Top50 signatures that connected with the CSC-specifically expressed genes

## Supplementary Figures legends

**Figure S1.**
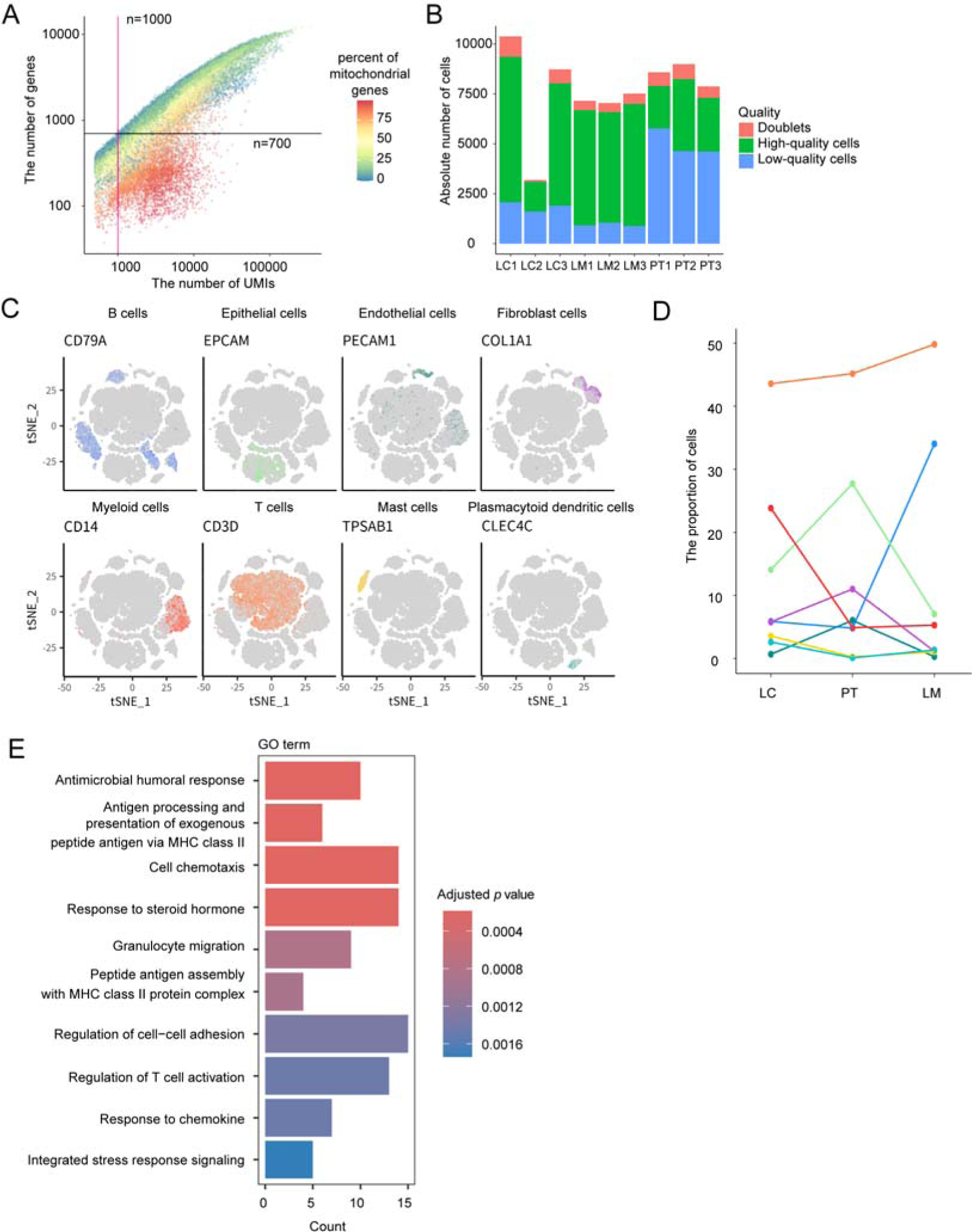
Single-cell RNA sequencing analysis in LSCC samples. **A.** Point plot shows the number of genes and unique molecular identifiers (UMIs) in each cell. Each dot represents a single cell; the color of each dot is according to the percentage of mitochondrial genes in the cell. **B.** The absolute number of cells from each patient, including doublets, high-quality cells, and low-quality cells. The high-quality cells were selected for downstream analysis. **C.** The t-SNE plot shows the authority marker genes expression in eight major cell types. **D.** Average proportions of different cell types in LC, LM, and PT. **E.** GO enrichment analysis of marker genes in LC-derived epithelial cells compared with PT-derived epithelial cells (adjusted *p* value < 0.01, log_2_FC > 1).

**Figure S2.**
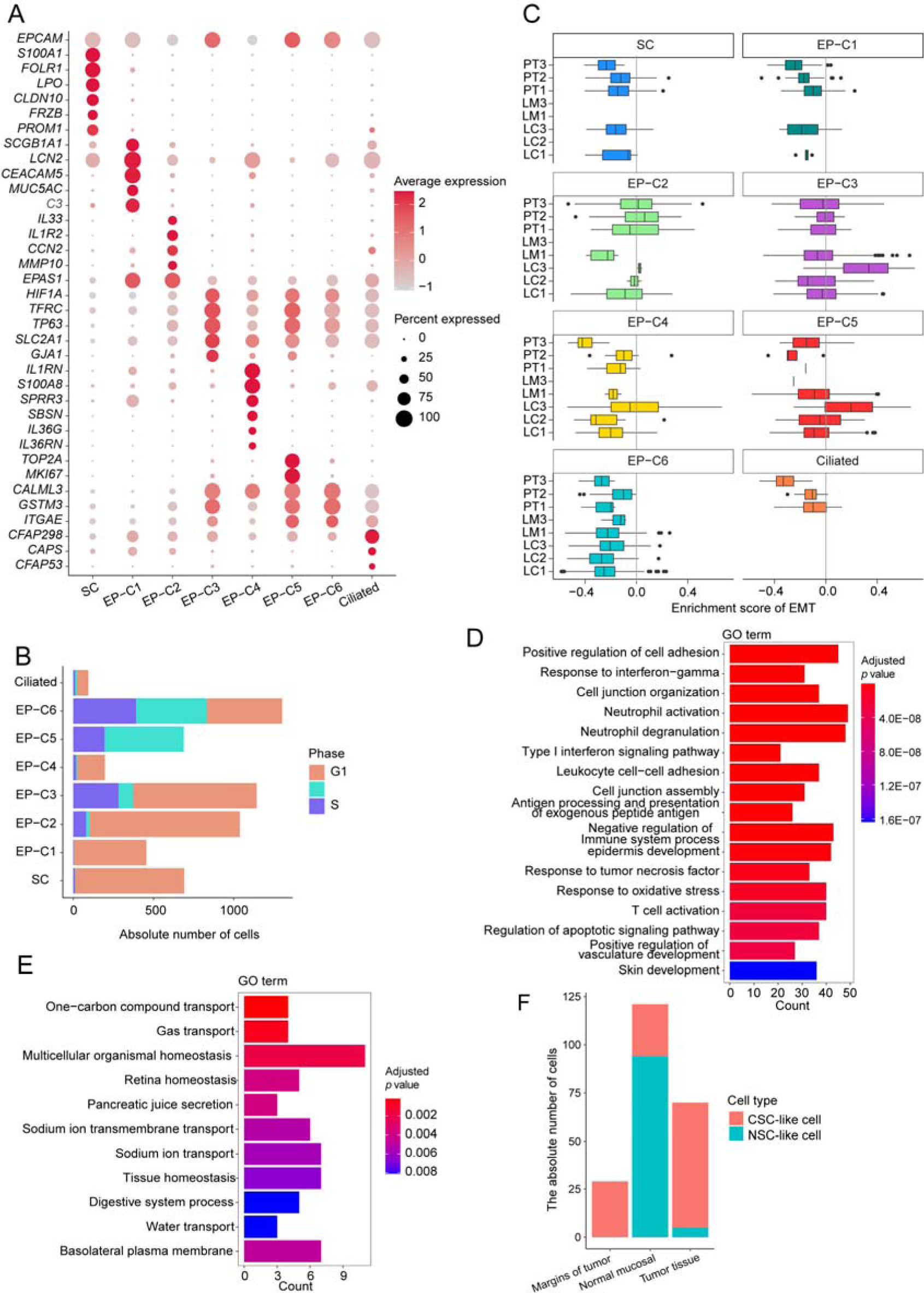
Characteristics of the epithelial cell subpopulations and cancer stem cell in LSCC. **A.** Dot plot shows the expression of marker genes in all the epithelial cell subpopulations. **B.** Bar plot shows absolute number of all the epithelial cell subpopulations in different cell cycle phases. **C.** GSVA scores of EMT gene sets in the epithelial cell subpopulations in LSCC. **D.** The enriched GO terms for marker genes in SC-C2. **E.** The enriched GO terms for marker genes in SC-C1. **F**. The distribution of CSC-like cells and NSC-like cells in different tissue type from an independent validation scRNA-seq dataset (GES206332).

**Figure S3.**
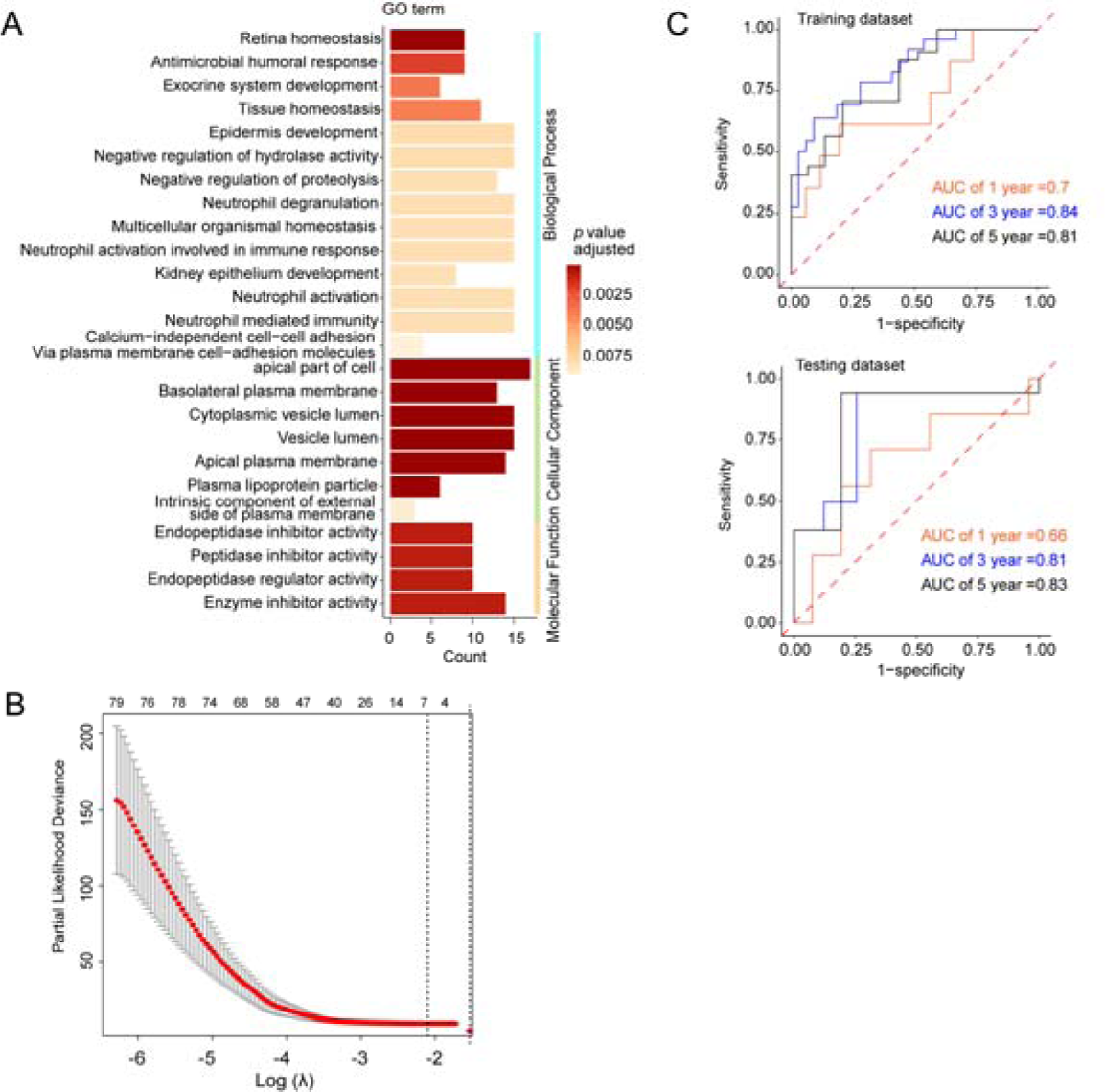
CSC marker gene signatures and clinical application in LSCC. **A.** Function enrichment analysis results for the CSC marker genes. **B.** The values of lambda used in the 10-fold cross-validation for glmnet. **C.** The ROC curves show predicted 1-, 3-, and 5-year overall survival of the LSCC patients based on prognostic model in the training and testing datasets.

**Figure S4.**
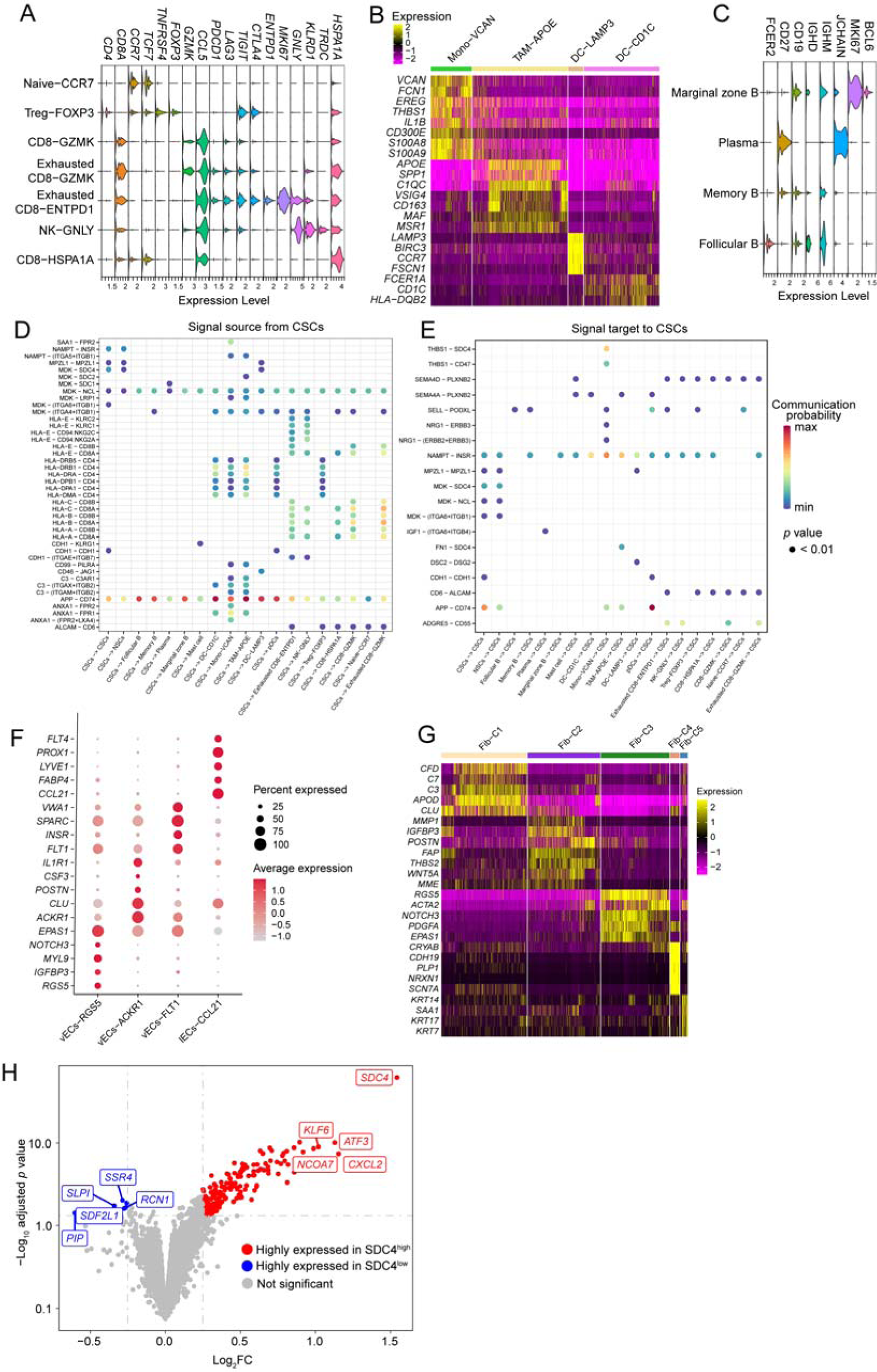
Analysis of the stroma cell subpopulations and The cell-cell communication between immune cells and CSCs in LSCC. **A.** Violin plot showed the conventional marker genes and differentially expressed genes in different subpopulations of T cells. **B.** Heat map showed the differentially expressed genes in different subpopulation of myeloid cells. **C.** Violin plot shows the conventional marker genes and differentially expressed genes in different subpopulation of B cells. **D**–**E.** Dot plot shows the significant ligand-receptor pairs between CSCs and immune cells. The dot color represents the calculated communication probability; dot size represents *p* values. **F.** Dot plot shows differentially expressed genes in different subpopulations of endothelial cells. **G.** Heat map shows differentially expressed genes in different subpopulations of fibroblast cells. **H.** Volcano plot shows differentially expressed genes between *SDC4*^high^ and *SDC4*^low^ groups of CSCs.

